# LARIS enables accurate and efficient ligand and receptor interaction analysis in spatial transcriptomics

**DOI:** 10.1101/2025.11.26.690796

**Authors:** Min Dai, Tivadar Török, Dawei Sun, Vallari Shende, Grace Wang, Yuesang Lin, Sherry Jingjing Wu, Alyssa Rukshin, Gord Fishell, Fei Chen

**Affiliations:** Stanley Center for Psychiatric Research, Broad Institute of MIT and Harvard, Cambridge, MA 02142, USA; Harvard Medical School, Blavatnik Institute, Department of Neurobiology, Boston, MA 02115, USA; Broad Institute of MIT and Harvard, Cambridge, MA 02142, USA; Department of Stem Cell and Regenerative Biology, Harvard University, Cambridge, MA 02138, USA

## Abstract

Advances in spatially resolved transcriptomics provide unprecedented opportunities to characterise intercellular communication pathways. However, robust and computationally efficient incorporation of spatial information into intercellular communication inference remains challenging. Here, we present LARIS (<u>L</u>igand <u>A</u>nd <u>R</u>eceptor Interaction analysis in <u>S</u>patial transcriptomics), an accurate and scalable method that identifies cell type-specific and spatially restricted ligand-receptor (LR) interactions at single-cell or bead resolution. LARIS is compatible with all spatial transcriptomic technologies and quantifies specificity, infers sender-receiver directionality, and detects how differential interactions vary across time and space. To compare LARIS with existing methods, we established a simulation framework to generate ground truth of LR interactions with defined tissue architecture and gene expression patterns. LARIS demonstrates superior performances over other methods in accuracy and scalability. We further applied LARIS to human tonsil and developing mouse cortex spatial transcriptomics datasets collected from various spatial techniques. This uncovered the signalling mechanisms shaping tissue organisation and their changes over time. LARIS reveals cell type-, niche-, and condition-specific signalling and scales to hundreds of thousands of cells in minutes. This provides an efficient and direct method for discovering the molecular interplay between apposed cells across development.

## Introduction

Intercellular communication orchestrates tissue development, homeostasis, and responses to stress and disease^1^. Computational inference of ligand-receptor (LR) interactions from transcriptomic data has therefore become a cornerstone for charting cellular interactions in complex tissues. However, computationally inference of cell-cell communication (CCC) faces challenges, including data sparsity, and spatial heterogeneity. Although recent advances in CCC algorithms have attempted to address some of these challenges by gene expression imputation and incorporating spatial constraints on LR interactions at populational level, they still struggle with low sensitivity and specificity, missing relevant interactions, producing many false-positive hits, respectively. Thus, sensitive, spatially resolved, and single-cell level reconstruction of intercellular communication networks remains a major unmet need^1^.

To meet these needs, we introduce LARIS (<u>L</u>igand <u>A</u>nd <u>R</u>eceptor Interaction analysis in <u>S</u>patial transcriptomics) as an accurate method for LR interaction and CCC inference in spatial transcriptomic data. LARIS enhances the spatial interaction detection by combining *in silico* LR diffusion and local cellular heterogeneity via a spatial *k*-nearest neighbours (*k*-NN) graph, to infer single-cell level LR interaction scores and spatial specific CCC with cosine similarity. To rigorously compare our success with previous efforts, we also establish a spatial transcriptomics simulation framework that provides a reference ground truth for benchmarking CCC methods. We show that LARIS outperforms existing methods, accurately recovering spatial and cell type-specific interactions while maintaining low spatial false-positive rates. We further demonstrate the capabilities of LARIS in two case studies: (i) a human tonsil Slide-tags dataset^2^, where LARIS resolves niche- and cell-state-specific communication and recovers both known and novel signals; and (ii) a developing mouse cerebral cortex Stereo-seq dataset^3–5^, where LARIS identifies layer-specific interactions and quantifies their temporal dynamics during development. Taken together, LARIS provides an accurate, scalable, and adaptable framework to infer ligand and receptor dynamics and decipher intercellular communication in complex biological systems.

## Results

### Overview of LARIS

LARIS is designed to identify and characterize LR interactions from spatial transcriptomics data using known databases and is compatible with both single-cell and bead/spot-based spatial datasets, enabling the discovery of cellular communication across diverse platforms (Fig. 1a and Extended Data Fig. 1). To improve both sensitivity and specificity, LARIS uses interaction abundance and cosine similarity to identify biologically relevant interactions, and a spatial *k*-NN graph to ensure that only spatially proximal cells are identified as interacting.

**Fig. 1:**
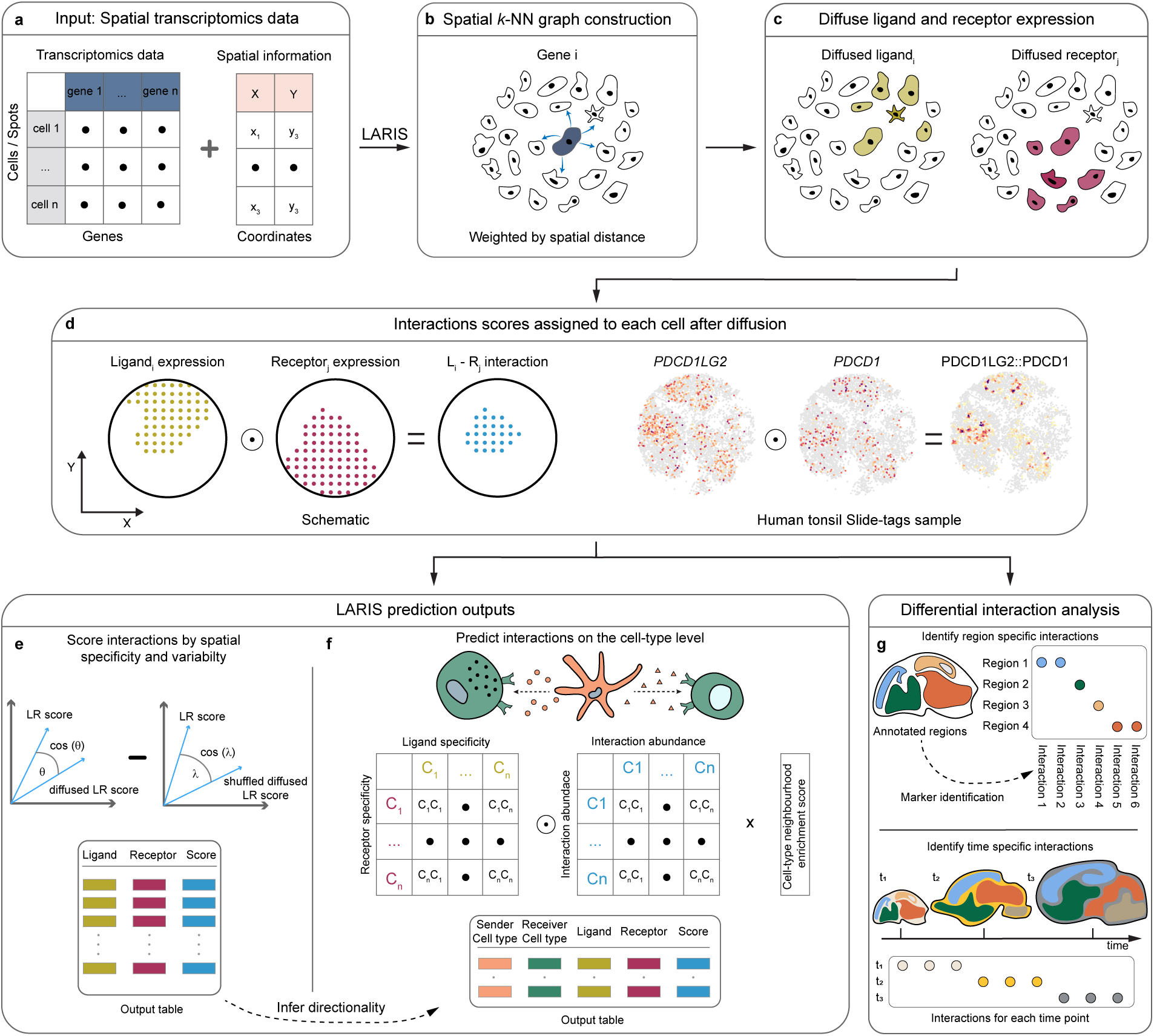
Overview of LARIS and its applications. **a,** Input data. LARIS relies on spatial transcriptomics data as input where each cell or spot is characterised by gene expression profiles with spatial coordinates (X, Y). **b,** A spatial *k*-nearest neighbour (*k*-NN) graph is constructed and weighted by the Euclidean distance between each cell or spot. **c,** In *silico* diffusion of ligand and receptor gene expression by the spatial *k*-NN graph. **d,** Calculate a ligand-receptor (LR) interaction score for each cell or spot by multiplying the diffused ligand and receptor expressions. For example, in the Slide-tags human tonsil sample, the gene expression of programmed cell death 1 ligand 2 (*PDCD1LG2*) and its receptor programmed cell death 1 (*PDCD1*) are used to calculate the interaction score PDCD1LG2::PDCD1 for each cell. **e,** Spatial specificity and variability is measured by calculating the cosine similarity (cos θ) between the observed and randomly shuffled diffused LR scores. Stronger similarity means more specific and stronger spatial localisation for an interaction. **f,** Cell type level interaction directionality is calculated by combining a specificity metric of the ligand and receptor, the abundance of LR scores in each cell type, and the enrichment of each cell type in each other’s neighbourhoods. **g,** Differentially enriched interactions can be identified by first labelling spatial regions, time points or conditions, then applying a differential test on the LR scores. For example, by labelling different regions of the tissue the differential interactions can be identified.

LARIS begins by constructing a *k*-NN graph weighted by the physical distance between cells or spots (Fig. 1b). Using this spatial graph, the normalized ligand and receptor expression values are spatially diffused, modelling the local availability of each (Fig. 1c). An LR interaction score is then calculated for each cell or spot by element-wise product (i.e., Hadamard product) of the diffused ligand and receptor expression (Fig. 1d). To prevent diffusion artifacts from generating false-positive interactions, this score is set to zero if the cell or spot did not express either the ligand or the receptor in the original gene expression data. This scoring at the individual-cell or spot level increases sensitivity for interactions involving rare cell types and reveals communication patterns within subpopulations that are otherwise obscured by cluster-level or aggregated methods (e.g., CellChat^6^ and CellPhoneDB^7^).

Building on these LR scores at single-cell/spot resolution, LARIS provides three complementary analytic approaches to discover the most specific interactions. First, LARIS identifies LR interactions with strong spatial patterns (Fig. 1e) using a cell type-agonistic approach. LARIS computes the cosine similarity between the calculated LR interaction scores and a spatially diffused version of those same scores. Building on the idea of negative sampling in network representation learning^8^, this observed similarity score is then corrected against a null value derived from spatially shuffled data. This yields a final spatial-specificity score for each LR pair (see Methods), of which the higher value indicates a stronger spatially localized pattern.

Second, LARIS infers communication directionality between sender and receiver cell types (Fig. 1f). To achieve this, it quantifies: (i) the cell-type specificity of ligand and receptor expression, (ii) the abundance of each LR pair for every cell type pairing, and (iii) a neighbourhood-enrichment score to detect co-localizing interactions between cell types. Combining this information with the LR spatial-specificity score yields a score for each LR pair in each sender-receiver cell type pair, which enables further prioritization of LR interactions based on cell types of interest and analysis of cell type communication patterns.

Third, LARIS identifies differentially active interactions among different conditions or contexts, such as predefined tissue regions/niches, developmental stages or disease progressions (Fig. 1g). LARIS facilitates statistical testing on the single-cell/spot level LR interaction score to discover LR pairs significantly enriched or depleted in one condition or context compared to another (see Methods).

The single-cell/spot level quantification of LR interaction strength, coupled with the three-views interpretation, i.e., spatial specificity, cell type directionality and differential interaction among conditions, enables LARIS to be an efficient and accurate method for characterizing the LR interaction and cell-cell communication dynamics.

### Benchmarking on simulated data

Prior work has highlighted low concordance among CCC methods, with major differences in top-ranked predictions^9^. These discrepancies stem from divergent LR databases, the inclusion of interactions lacking experimental validation, and distinct model architectures^1,9^. A persistent challenge in the field has been the benchmarking of these tools, which is hampered by the absence of comprehensive experimentally validated ground-truth datasets.

To address this gap, we designed a simulation framework to generate spatial transcriptomics data with real-world tissue characteristics and known ground-truth LR interactions, providing full control over cell locations, gene expression (including genes that encode ligands and receptors), and active LR pairs with different levels of spatial and cell type specificities (Fig. 2a). This methodology enables rigorous, quantitative performance comparisons.

**Fig. 2:**
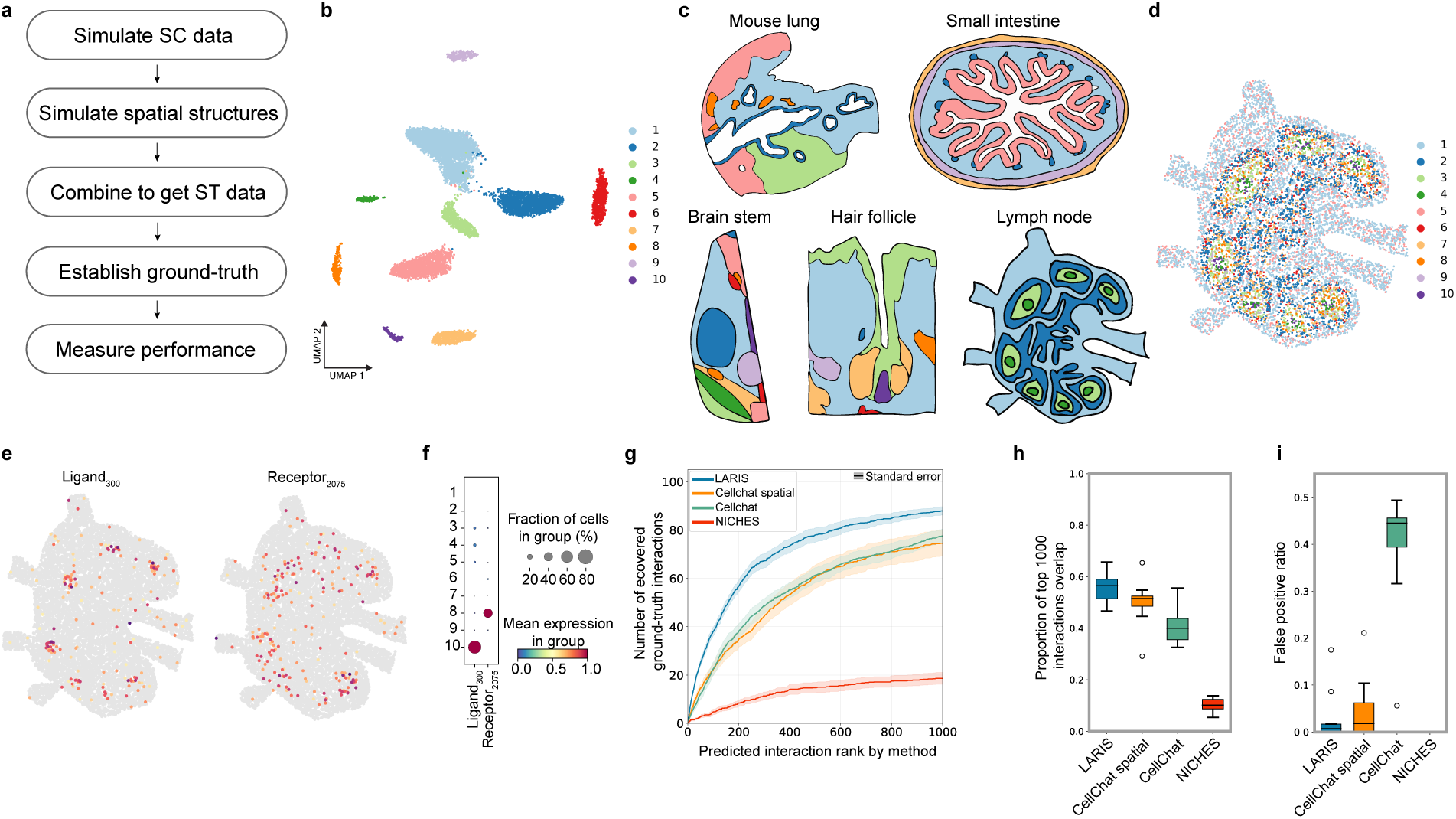
Benchmarking on simulated data. **a,** Simulated spatial transcriptomic based cell-cell communication benchmarking workflow. **b,** Uniform manifold approximation and projection (UMAP) of simulated scRNA-seq data coloured by cluster annotations. **c,** Simulated spatial structures based on real tissue examples, such as infected mouse lung, small intestine, developing brain stem, hair follicle, and lymph node. **d,** Combined scRNA-seq data with the spatial structure for the lymph node example. **e,** Spatial gene expression mapping of a simulated, interacting ligand-receptor pair for Ligand_300_ and Receptor_2075_.Ground-truth interaction is between simulated cluster 10 (sender) and cluster 8 (receiver). **f,** Expression dot plot of simulated Ligand_300_ and Receptor_2075_. **g-i**, Performance metrics between the ground-truth and LARIS, CellChat spatial, Cellchat, and NICHES prediction outputs. **g)** Recovery of the number of top 100 ground-truth interactions by method. **h)** Pairwise overlap with ground-truth among the top 1,000 predicted interactions across methods. **i)** False positive rates of predicted interactions across methods.

The workflow first simulates scRNA-seq data using Splatter^10,11^. Simulation parameters were calibrated to ensure that cell type identities are distinct and that quality metrics—such as dropout rates, unique molecule identifier (UMI) distributions, and gene variance—closely mimic those of a real-world reference, a human tonsil Slide-tags single nuclei RNA-seq dataset^2^ (Fig. 2b and Extended Data Fig.2). Next, these simulated cells are mapped onto complex spatial structures, which are generated either *de novo* using scCube^12^ or by templating from real tissue architectures, including mouse lung, small intestine, and developing brainstem, hair follicle and lymph node (Fig. 2c, d).

To establish a ranked ground truth, we defined an interaction’s importance based on its expression specificity and spatial likelihood. First, we computed a ‘relevance score’ for each ligand and receptor in different cell types based on the simulation parameters that account for its simulated expression abundance and cell-type specificity (Extended Data Fig.3; see Methods). The ground-truth LR interaction ranking was then generated by multiplying these ligand and receptor relevance scores across all cell-type pairings. This comprehensive list was subsequently filtered for spatial feasibility, resulting in a final, ranked set of active LR interactions with varying specificities, known to occur between spatially proximal cell types (Fig. 2e, f).

**Fig. 3:**
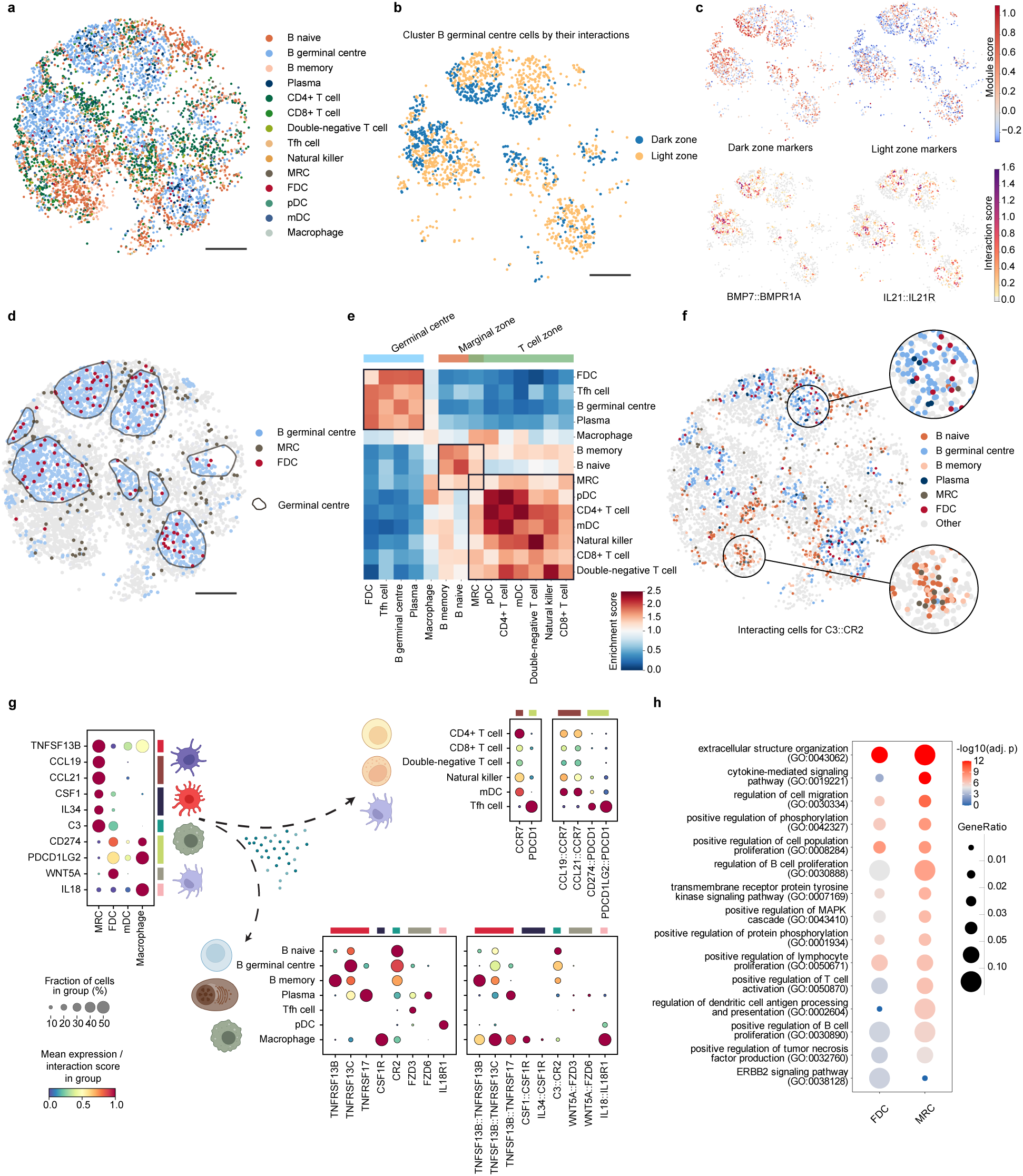
LARIS quantifies cellular communication on the single-cell level in the human tonsil. **a)** Spatial mapping of snRNA-seq profiles, coloured by cell type annotations. Tfh cell, T follicular helper cells; MRC, marginal reticular cells; FDC, follicular dendritic cells; pDC, plasmacytoid dendritic cells; mDC, myeloid dendritic cells. **b)** Clustering of B germinal centre cells by their interaction scores separate their dark and light zone identity. **c)** Gene expression modules of known dark-zone genes (CXCR4, AICDA, FOXP1, NME) and light-zone genes (CD83, LMO2, BCL2A1), along with the interactions BMP7::BMPR1A (dark zone) and IL21::IL21R (light zone), which are enriched in their respective zones. **d)** Spatial mapping of B germinal centre, MRC, and FDC cells coloured as in (**a)**, rest of the cells are coloured in grey. Germinal centres indicated by the grey outlines. **e)** Spatial neighbourhood enrichment scores by cell type label. A value of 1 indicates random expectation; values >1 (brown) indicates enrichment and values <1 (blue) indicate depletion. Spatial regions indicated by the black boxes. **f)** The complement system is mediated by MRC and FDC cells at different locations of the tonsil as both express C3. Interacting cells coloured and annotated as in (**a).** MRCs primarily interact with B naïve, B memory, while FDCs primarily interact with B germinal centre and Plasma cells. **g)** Dot plots of selected ligand genes (TNFSF13B, CCL19, CCL21, CSF1, IL34, C3, CD274, PDCD1LG2, WNT5A, IL18), receptor genes (CCR7, PDCD1, TNFRSF13B, TNFRSF13C, TNFRSF17, CSF1R, CR2, FZD3, FZD6, IL18R1), and corresponding LR interaction scores across the cell types participating in these interactions. MRCs act as key signalling hubs as they influence almost all cell types in the tonsil. **h)** Gene ontology of biological processes between MRC and FDC populations based on their differential communication. Scale bars, 500 μm.

Using this ground-truth simulation framework, we compared LARIS against other CCC methods, including CellChat (non-spatial)^13^, CellChat-spatial (v2)^6^, and NICHES^14^. First, we assessed the efficiency of each method in prioritizing the top ranked ground-truth LR interactions. We plotted recovery curves, showing the number of top ground-truth hits recovered as a function of increasing prediction rank (Fig. 2h and Methods). LARIS demonstrated a higher area under the curve, indicating it successfully ranked the most critical ground-truth interactions higher than all other methods (Supplementary Table 1). The non-spatial and spatial versions of CellChat performed similarly, while NICHES recovered substantially fewer of these high-priority interactions. We next evaluated the overall recall by calculating the proportion of ground-truth interactions recovered within the top 1,000 predictions of each method (Fig. 2h). LARIS again performed best, and CellChat-spatial outperformed the original CellChat, confirming that the incorporation of spatial information improves discovery. Finally, we evaluated the spatial false-positive rate, defined as the proportion of predicted interactions occurring between cell types that were not spatially proximal in the simulation. LARIS and NICHES both exhibited very low false-positive rates (Fig. 2i). In contrast, CellChat performed worst, with over 40% of its predictions being spatially false positives, highlighting the critical advantage of integrating spatial information for CCC inference.

Taken together, our spatial simulation framework provides a unified and unbiased approach for benchmarking CCC methods’ performance on spatial transcriptomics data. Across all metrics, LARIS outperformed existing methods, demonstrating superior recovery of spatial and cell type-specific, high-priority interactions while maintaining a low spatial false-positive rate, thereby validating the efficacy of our approach.

### LARIS identifies ligand-receptor interactions defining zonations of germinal centres in human tonsil

We next tested LARIS’s ability to profile cellular communications on the human tonsil Slide-tags dataset^2^, as this enables us to assess/quantify LR dynamics at single-nucleus resolution within a tissue with well-characterised spatial structures and LR interactions, offering a robust reference for benchmarking (Fig. 3a and Extended Data Fig. 4a)^15^.

First, we asked whether LR scores alone could distinguish dark- and light-zone identity in germinal centre (GC) B cells, a task that is challenging with scRNA-seq. The dark zone hosts proliferating B cells undergoing somatic hypermutation; the light zone mediates affinity-based selection via follicular dendritic cells (FDCs) and T follicular helper (Tfh) cells^15^. *De novo* clustering of LR scores separated GC B cells (Fig. 3b and Extended Data Fig. 4b), a pattern validated by the expression of canonical dark zone and light zone gene modules^16^ (Fig. 3c). Zone-specific enrichment analysis revealed distinct communication programs: *BMP7* and its receptors interact in the dark zone; *BMP7* signalling has been reported to induce apoptosis and negatively regulate GC B-cell survival^17,18^. In contrast, *L21-IL21R* interactions, characteristic of Tfh-derived signalling, were enriched in the light zone, supporting the role of IL21 in promoting GC B-cell expansion and early plasma cell differentiation^19,20^ (Fig. 3c and Extended Data Fig. 4c, d).

### LARIS identifies marginal reticular cells as key signalling hubs in human tonsil

Reclustering using the LR scores combined with gene expression separated FDCs from marginal reticular cells (MRCs), revealing two distinct signalling patterns of cell populations within the originally annotated FDCs (Extended Data Fig. 4e). The identities of FDCs and MRCs were validated using canonical marker genes (Extended Data Fig. 4f)^21^, and spatial mapping confirmed their respective localization, with FDCs restricted to germinal centres and MRCs positioned in the surrounding regions (Fig. 3d).

To further characterise regional niches, we computed neighbourhood enrichment scores for each cell type (Fig. 3e). Unsupervised clustering on cell neighbourhood enrichment has clearly separated the germinal centres, B, and T cell rich zones. FDCs primarily communicate with germinal-centre cells, including Tfh and GC B, and plasma cells, whereas MRCs interact more broadly with naïve and memory B cells as well as pDC, mDC, and multiple T-cell subsets. Interestingly, macrophages showed no strong enrichment in any of those regions, contributing to regulation in all regions of the tonsil^22^.

While FDCs and MRCs occupy distinct niches and differ in their interacting partners, they converge in specific signalling programs. Both express C3, a key complement component involved in antigen presentation^23,24^ (Fig. 3f), and MRCs additionally produce soluble regulators of the classical, lectin, and alternative pathways^25–28^ (Extended Data Fig. 4j).

We also discovered distinct LR interactions associated with FDCs and MRCs: Inside the germinal centres, FDCs and macrophages express the checkpoint ligands PD-L1 (CD274) and PD-L2 (PDCD1LG2), positioning them to modulate Tfh cells through PD-1 (PDCD1)^29^. FDCs also uniquely express WNT5A, which supports germinal-centre B-cell survival via non-canonical WNT signalling^30^; in our data, we identified FZD3 as the likely interacting partner. Interestingly, we found that GC B cells express high levels of *PDGFD*, which could influence the clonal expansion and differentiation of the *PDGFRβ* positive MRCs into FDCs^31–34^ (Extended Data Fig. 4g,h).

Outside the germinal centres, MRCs express TNFSF13B (BAFF), often considered a defining FDC product^35,36^. We found that FDCs within the germinal centres express very low levels (Fig. 3e), consistent with flow cytometry and scRNA-seq data^21^. TNFSF13B is an essential survival factor for naïve B cells^35^, although it is dispensable for germinal-centre B-cell survival^37^. In our dataset, macrophages and mDCs were also substantial sources of TNFSF13B, supporting their contribution to B-cell survival and differentiation. Lack of TNFSF13B leads to failure of FDC maturation; consequently, stimulation of B cells via immune complexes can not occur^38^. Beyond their effects on B cells, MRCs help organise T cell-rich zones in the paracortex by secreting CCL19 and CCL21, chemokines that recruit CCR7-positive cells^39^, including T cells, NK cells, and mDCs in the tonsil. MRCs supply CSF1 and IL34 which are important growth factors required for the control of monocyte and macrophage differentiation, survival, proliferation, and renewal^40^, suggesting their importance in macrophage development and differentiation outside of germinal centres. These findings highlight the importance of MRCs as tissue organisers in tonsil that signal to B cells, T cells, and macrophages shaping their development, survival, and migration (Extended Data Fig. 4i).

To compare pathway-level communication between MRCs and FDCs, we performed a differential analysis of LR interactions between the two populations. The Gene Ontology (GO) analysis revealed both shared and distinct biological processes (Fig. 3h)^41^: For example, GO terms related to dendritic cell antigen processing and presentation were significantly enriched in MRCs, whereas the ERBB2 signalling pathway was enriched in FDCs.

By applying LARIS to a human tonsil Slide-tags sample, we demonstrated that it can resolve distinct cell states within a cell type and identify the LR interactions enriched for those states. Its single-cell level scoring further guided the analysis by revealing spatially distinct communication patterns of FDCs and MRCs. LARIS recovered known LR pairs and connected them to sender and receiver cell types in the tonsil and suggested previously uncharacterised communications that influence cell survival, migration, and maturation.

### LARIS identifies region and time specific interactions in the developing mouse brain

We next applied LARIS to map the spatiotemporal dynamics of LR interactions during mouse brain development, a process governed by spatially organized signalling cascades^42^. We focused on the cerebral cortex, a structure defined by its distinct layered architecture and corresponding functional and transcriptional heterogeneity^43^.

To assess this we applied LARIS to Stereo-seq datasets collected at different developmental stages of the mouse brain, i.e., embryonic day (E)14.5, E16.5, postnatal day (P)7, P14, and P77^3–5^ (Fig. 4a and Extended Data Fig. 5a). We annotated the cortex as upper and deep layers for the postnatal samples, and for the prenatal samples we included additional developmental cell types (Fig. 4a).

**Fig. 4:**
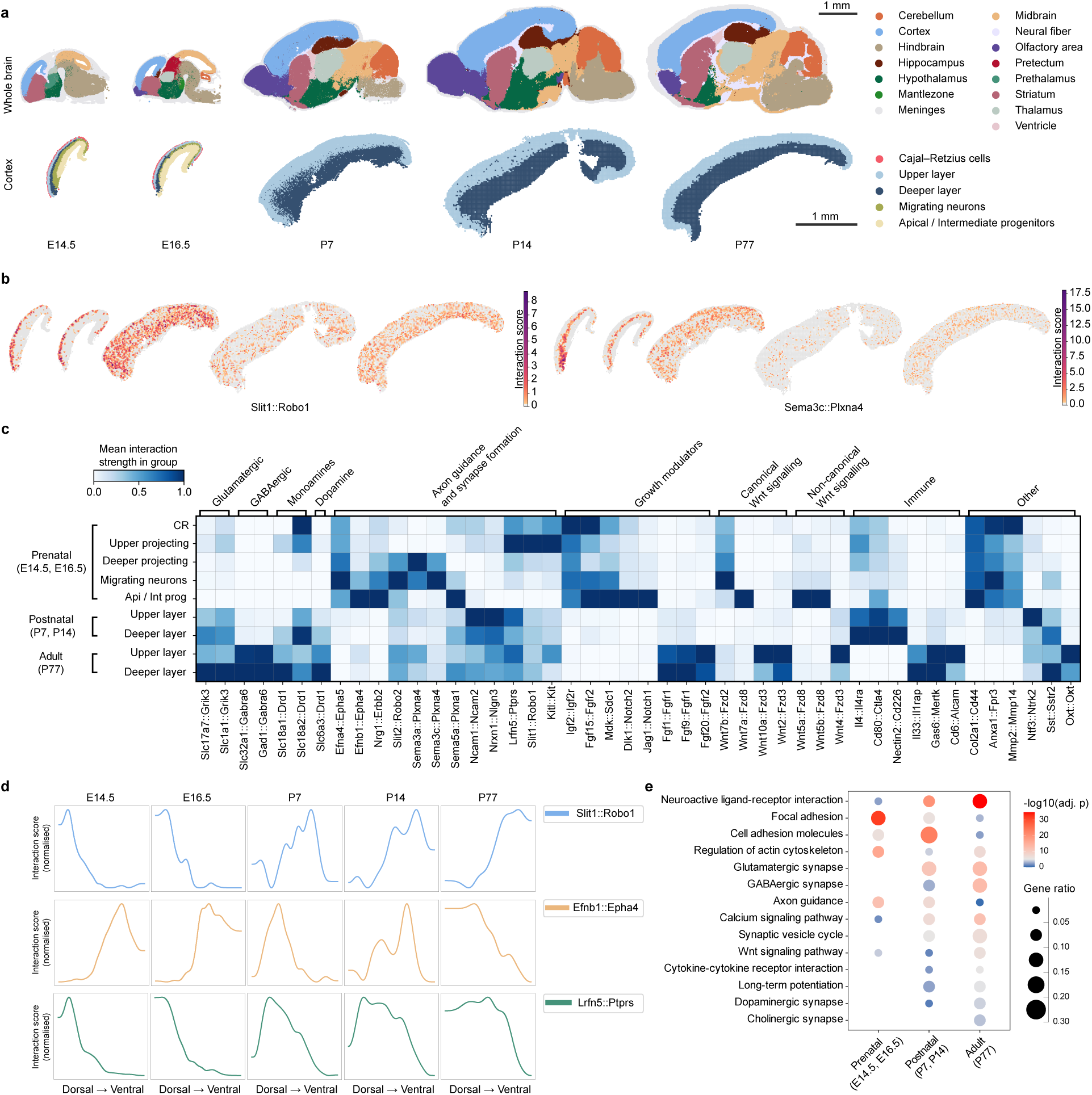
Spatiotemporal analysis of ligand-receptor interactions in the mouse brain cortex with LARIS. **a)** Spatial mapping of neuroanatomical and cortical regions profiled across prenatal (E14.5, E16.5), early postnatal (P7, P14), and adult (P77) stages. **b)** Slit1 and Robo1 interaction is highly specific to the upper layer projecting neurons and CR cells at the prenatal stage, then sporadically present across all layers. Sema3c and Plxna4 interaction is strong in the migrating and deep layer projecting neurons in the prenatal stage but then follows a similar trend and loses its spatial specificity after birth. **c)** Interacting ligand-receptor (LR) pairs in the cortical regions at different developmental time points. Cortical annotation classes include Cajal-Retzius (CR) cells, upper- and deep-layer, migrating neurons, and apical/intermediate progenitors (Api/Int prog). Interactions are grouped according to their functionality, such as neurotransmission (glutamatergic, GABAergic, monoamines, dopamine), axon guidance and synapse formation, growth modulators, canonical and non-canonical Wnt signalling, immune and other. Interaction scores are normalized per interaction. **d)** Changes in normalised LR interaction scores across the dorsal-ventral developmental axis at the different developmental time points for the Slit1::Robo1, Efnb1::Epha4, and Lrfn5::Ptprs interactions. **e)** KEGG pathway-level enrichment of interacting gene sets across stages (prenatal, postnatal, adult). Bubble size denotes gene ratio; colour indicates −log10(adjusted *P* value).

LARIS identified differentially enriched spatial LR interactions across mouse cortex development. For example, the Slit1::Robo1 interaction, known to modulates proliferation and neurogenesis^44,45^, axon guidance and neuronal migration^46,47^, was restricted to upper layer neurons prenatally but present in both upper and deep layer postnatally (Fig. 4b). Another intriguing example, the Sema3c::Plxna4 interaction, which was most prominent in the migrating and deep layer neurons prenatally, yet only sporadically present in the postnatal cortex (Fig. 4b). *Sema3a*, a closely related ligand with similar structure and expression to *Sema3c*, regulates interneuron positioning via *Plxna4*^48^. By contrast, the role of Sema3c in the developing mouse cortex remains unknown, raising the possibility of a Sema3c::Plxna4 signalling axis.

To systematically analyse the LR interactions identified by LARIS, we grouped them by function (e.g., neurotransmitters, axon guidance, growth modulators, Wnt signalling) and plotted their strengths across developmental stages and cortical layers (Fig. 4c). This revealed clear stage-specific activities patterns of different ligand-receptor pairs. For example, among the growth modulators, Igf2::Igfr2^49^, Fgf15::Fgfr2^50^, Mdk::Sdc1^51^, Dlk1::Notch2^52^, Jag1::Notch^53^ were primarily enriched prenatally In contrast, Fgf1::Fgfr1, Fgf9::Fgfr1, Fgf20::Fgfr2, were predominant in the adult brain, likely contributing to neuronal maintenance^54,55^.

To connect these LR interactions to underlying biological processes, we performed Gene Set Enrichment Analysis (GSEA)^41^ on the differentially expressed LR pairs at different stages (Fig. 4d). The results confirmed expected developmental trajectories: prenatal stages were enriched for gene function terms such as ‘axon guidance’, while the adult stage was dominated by pathways related to synaptic integration (e.g., ‘neuroactive ligand-receptor interaction’, ‘glutamatergic synapse’, ‘GABAergic synapse’, and ‘synaptic vesicle cycle’).

We also examined LR interactions along specific spatial axes. Along the dorsal-ventral axis, Slit1:Robo1, Efnb1::Epha4, and Lrfn5::Ptprs showed clear spatiotemporal dynamics (Fig. 4d). Along the rostral-caudal axis, we further identified multiple spatially restricted interactions (Extended Data Fig. 5b, c). For example, *Aldh1a3*, which encodes an enzyme that converts retinaldehyde into retinoic acid, was expressed at P7 specially in the caudal upper layer corresponding to the forming visual area. This pattern was confirmed by Allen Institute’s *in situ* hybridization (Extended Data Fig. 5d). Other examples include Cdh12:Itgb1 enrichment in the rostral cortex and *Penk*, an opioid polypeptide hormone, enrichment in the rostral deep layers postnatally (Extended Data Fig. 5b, c).

Together, these spatiotemporal analyses on mouse cortex development demonstrated that LARIS can identify and characterise key signalling molecules and pathways underlying the complex intracellular and intercellular dynamics through development.

## Discussion

Here we presented LARIS, a computational tool that allows the quantification of cell type and spatial LR interactions across time and space. We established a simulation-based ground-truth framework to measure and quantify its performance and demonstrated LARIS’s superior performance over existing methods. Applied to a human tonsil Slide-tags dataset, LARIS recovered canonical interactions underlying tissue organisation, cell migration, and immune responses, revealed previously unrecognised ligand sources, and separated cell states based on LR interaction patterns. In a Stereo-seq series of the developing mouse brain, LARIS resolved the spatial dynamics of LR interactions across cortical layers over multiple time points and linked these patterns to pathways central to brain development.

Computational inferences of cell-cell interactions are powerful for hypothesis generation but ultimately require experimental validation to establish mechanism. Incorporating spatial context reduces predicted false positives; however, co-localisation (even physical proximity) does not ensure a functional effect on receiver cells because other mechanisms might mask or complement active signalling through parallel signals. A further limitation is that transcriptomics measures mRNA, whereas proteins (and their post-translational modifications) drive signalling. Coupling to proteomic assays and integrating chromatin accessibility (e.g., ATAC-seq) to read out responses in receiver cells will strengthen causal interpretation.

Our benchmarking framework was designed to mimic real-world datasets and prioritise the reliable recovery of ligand-receptor (LR) pairs. A current limitation is that it does not directly assess downstream consequences of LR interaction, such as the activation of subsequent signalling or ultimately transcriptional programs by the receptors in the receiver cell types. The LR interaction scores generated by LARIS, however, does provide a quantitative foundation to bridge this gap; these scores could be correlated with differential gene expression in the receiver cell types to link LR interaction changes with functional effects. Future simulation frameworks that explicitly model the propagation of signalling cascades and downstream transcriptional changes would provide an invaluable reference for robustly evaluating such downstream predictions.

Many interactions involve co-receptors or multimeric ligands that act as complexes. LARIS does not yet model complex stoichiometry explicitly, which is an avenue for improvement. Given the zero-inflated nature of spatial assays, naively requiring all subunits to be co-detected would discard biologically plausible events lost to dropout and increase noise. Probabilistic complex scoring or partial-evidence models may offer a better balance between specificity and sensitivity.

We have demonstrated that LARIS can comparatively analyse and quantify interactions across time points; it could likewise be used to contrast signalling between conditions and matched controls, enabling differential LR communication analyses.

New spatial technologies are being developed constantly, and the field is moving towards the spatial profiling of cells compared to just dissociated single-cell data. LARIS will be an ideal tool to uncover and disentangle the hidden information inside the data and to point the most relevant focuses for future experimental planning. As the field moves towards better resolution and wider gene coverage, LARIS will likely be able to identify the most relevant ligand and receptors in the profiled sample and be ideal to analyse and quantify interactions across time and space.

## Extended Data Figures

**Extended Data Fig. 1:**
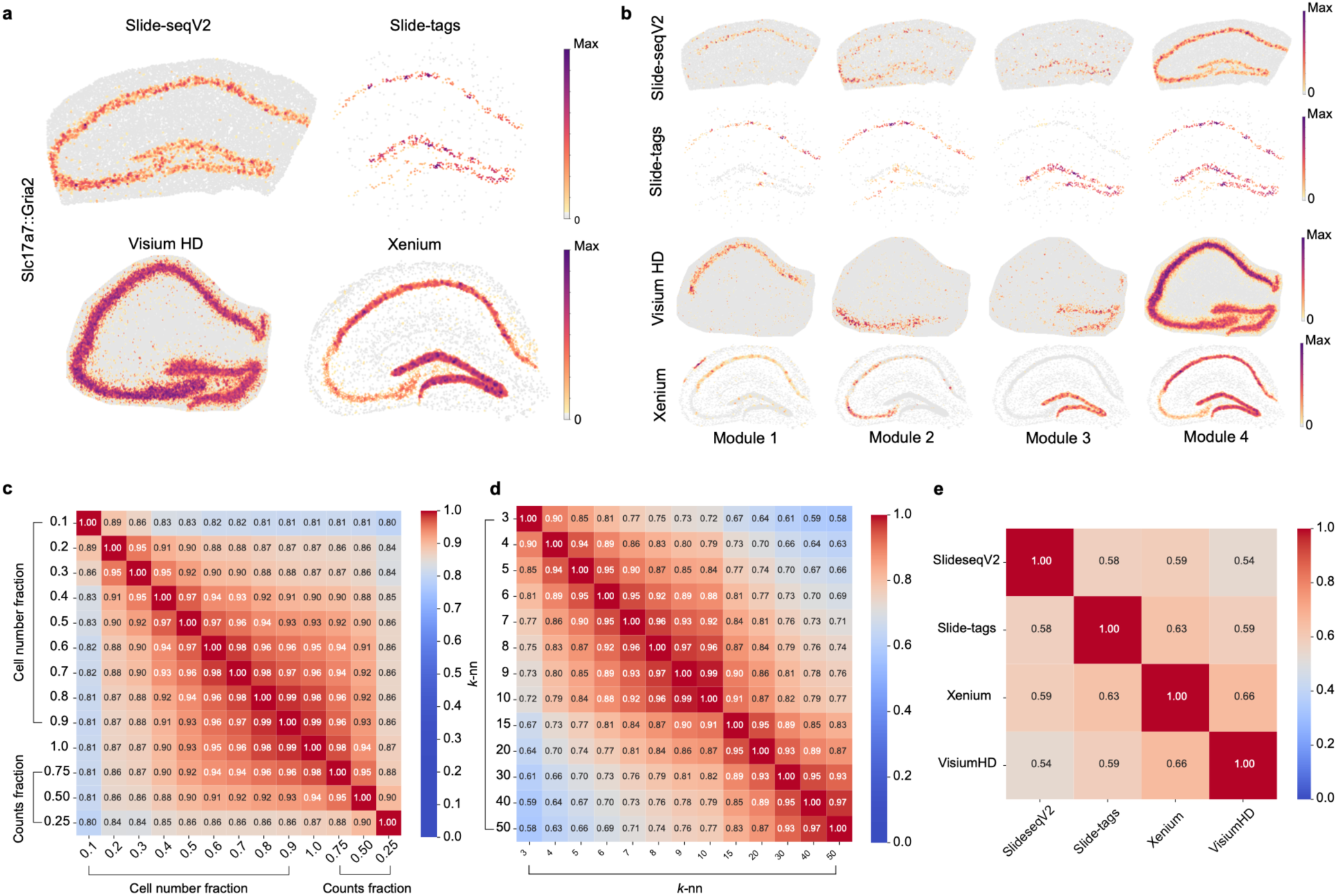
LARIS offers robust and comparable ligand-receptor interaction predictions from Slide-tags, Slide-seqV2, Visium HD, and Xenium datasets in the mouse hippocampus. **a)** LARIS is compatible with imaging, spot, and sequencing based spatial technologies. **b)** Spatial modules of ligand-receptor (LR) interactions in the mouse hippocampus. **c)** Spatially variability scores of LR interactions are robust when either the number of cells or detected genes counts is reduced. Shown are Pearson correlation coefficients between scores from the full dataset and downsampled subsets across downsampling levels. Xenium data of mouse brain hippocampus. **d)** Effect of various *k*-nn choices on spatially variable LR scores measured by Pearson correlation coefficients. Slide-tags data of human tonsil. **e)** Spatial variability scores for LR interactions are robust across spatial technologies. Shown are pairwise Pearson correlation coefficients between each method. All samples are mouse hippocampus.

**Extended Data Fig. 2:**
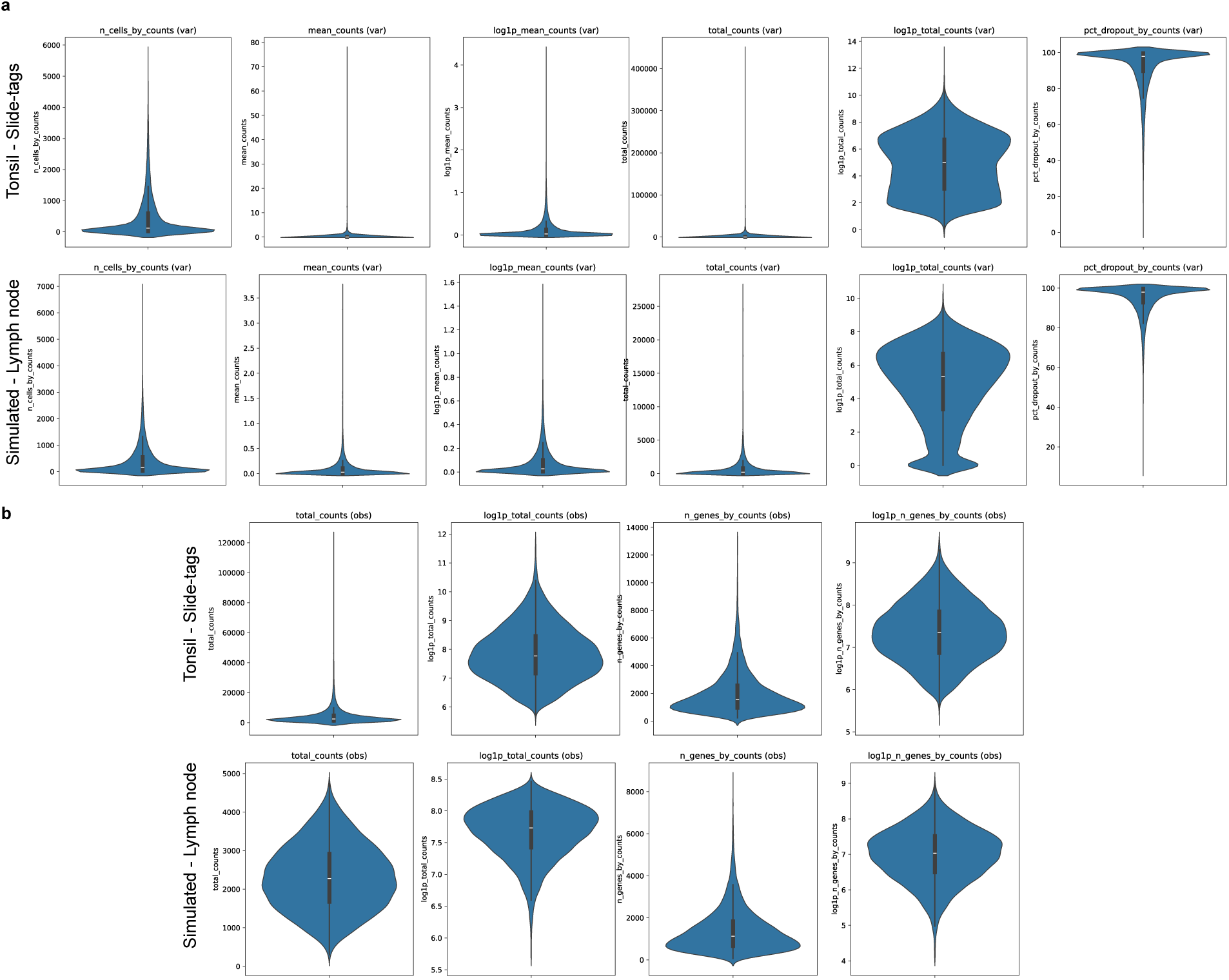
The simulated single-cell RNA-seq data accurately recapitulates the real life example of the human tonsil Slide-tags single nucleus RNA-seq data. **a)** Gene quality metrics. **b)** Cell quality metrics.

**Extended Data Fig. 3:**
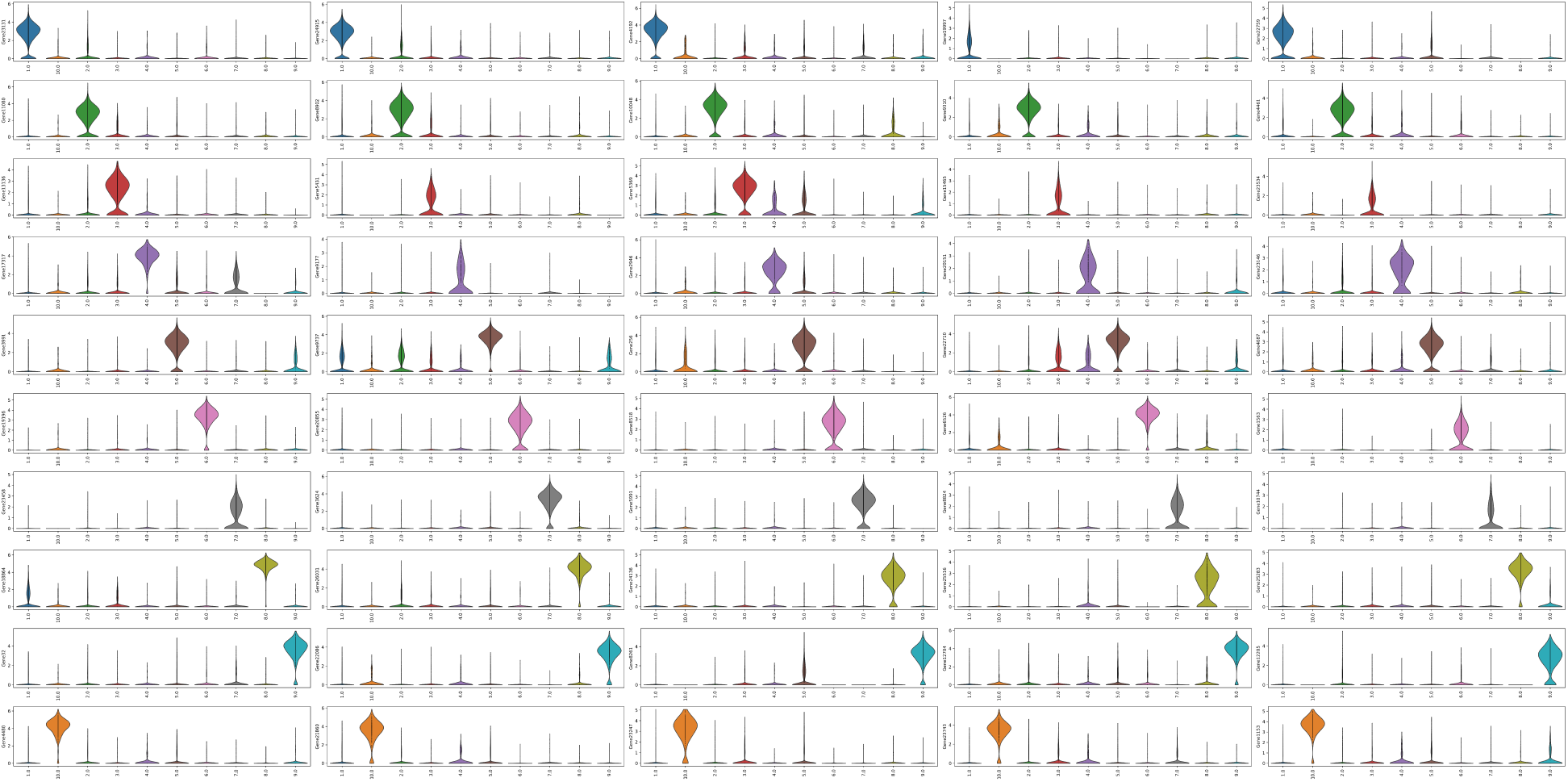
Top five genes by their relevancy scores for each simulated cluster based on their abundance and specificity for the simulated lymph node example.

**Extended Data Fig. 4:**
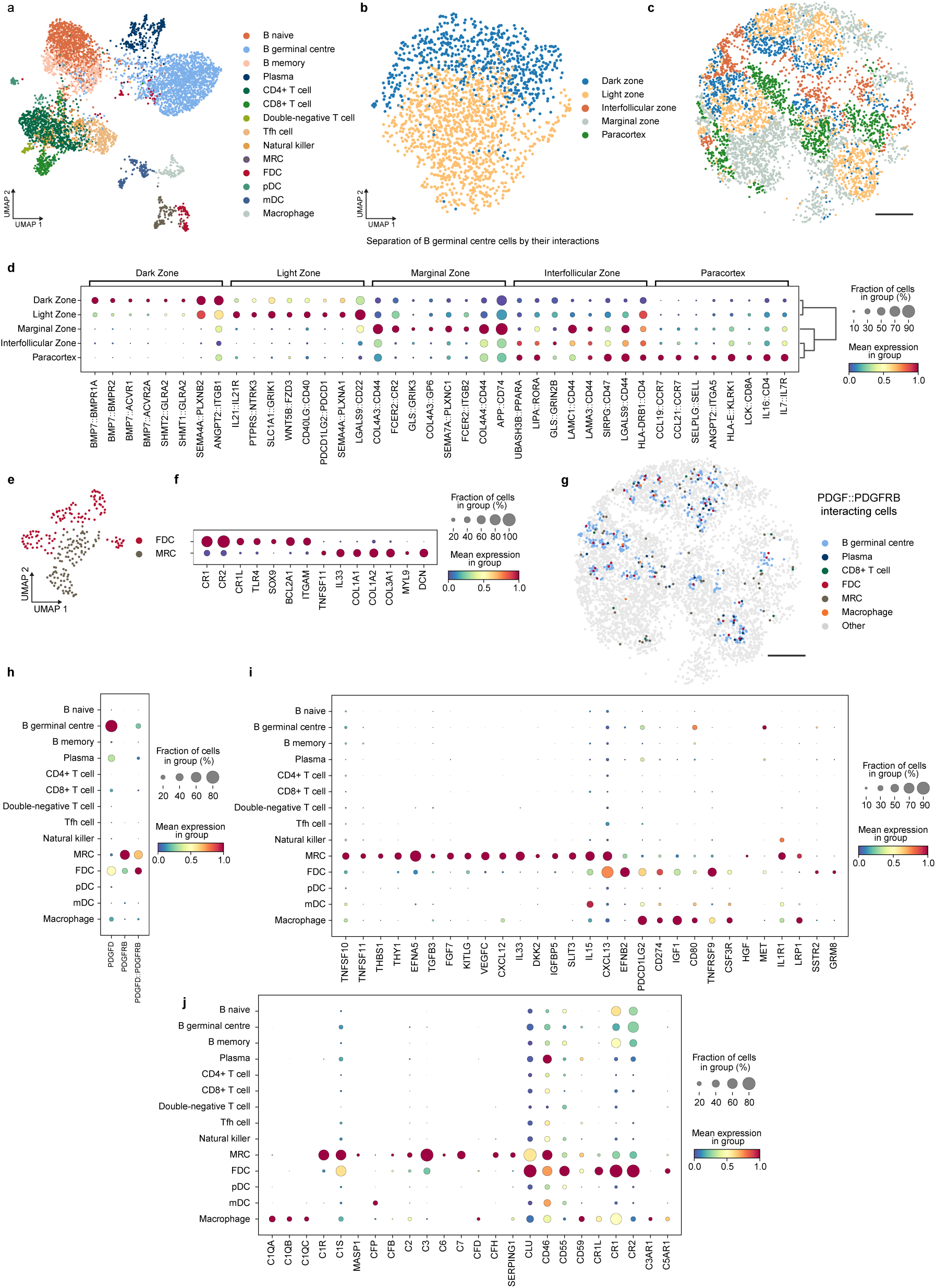
LARIS quantifies cellular communication on the single-cell level in the human tonsil. **a)** UMAP embedding of snRNA-seq profiles coloured by cell type annotations. Tfh cells, T follicular helper cells; MRC, marginal reticular cells; FDC, follicular dendritic cells; pDC, plasmacytoid dendritic cells; mDC, myeloid dendritic cells. **b)** Clustering of B germinal centre cells by their interaction scores separate their dark and light zone identity. UMAP embedding. **c)** Spatial mapping of functional regions in the tonsil. **d)** Differentially enriched ligand-receptor interactions in the functional regions of the tonsil reveal known ligands and receptors. BMP7 and its receptors in the dark zone promote B germinal center cell death, *Il21* and and its receptor *IL21R* regulate B germinal center maintenance, and *CCl19*, and *CCL21* and its receptor *CCR7* known as an important T cell homing signal in the paracortex. **e)** UMAP embedding of snRNA-seq profiles combined with LR interaction scores for the FDC and MRC populations. **f)** Dot plot of known marker FDC and MRC gene expression used to separate the two clusters. **g)** Platelet-derived growth factor D (PDGFD) is abundantly expressed by the B germinal centre cells, potentially triggering and contributing to the transition of MRCs into FDCs through the platelet-derived growth factor receptor beta (PDGFRB). Spatial mapping of interacting cells coloured as in **(a)**. **h)** Expression dot plot of PDGFD and PDGFRB and their LR scores. **i)** Multiple relevant ligand and receptor genes are specific for either the MRC, FDC, mDC, or macrophage populations. **j)** Complement system related genes. MRCs are likely important mediators of all three complement pathways, expressing multiple classical (C1R, C1S), lectin (MASP1), central components (C2, C3), and soluble regulators (CFB, CFH, SERPING1). Scale bars, 500 μm

**Extended Data Fig. 5:**
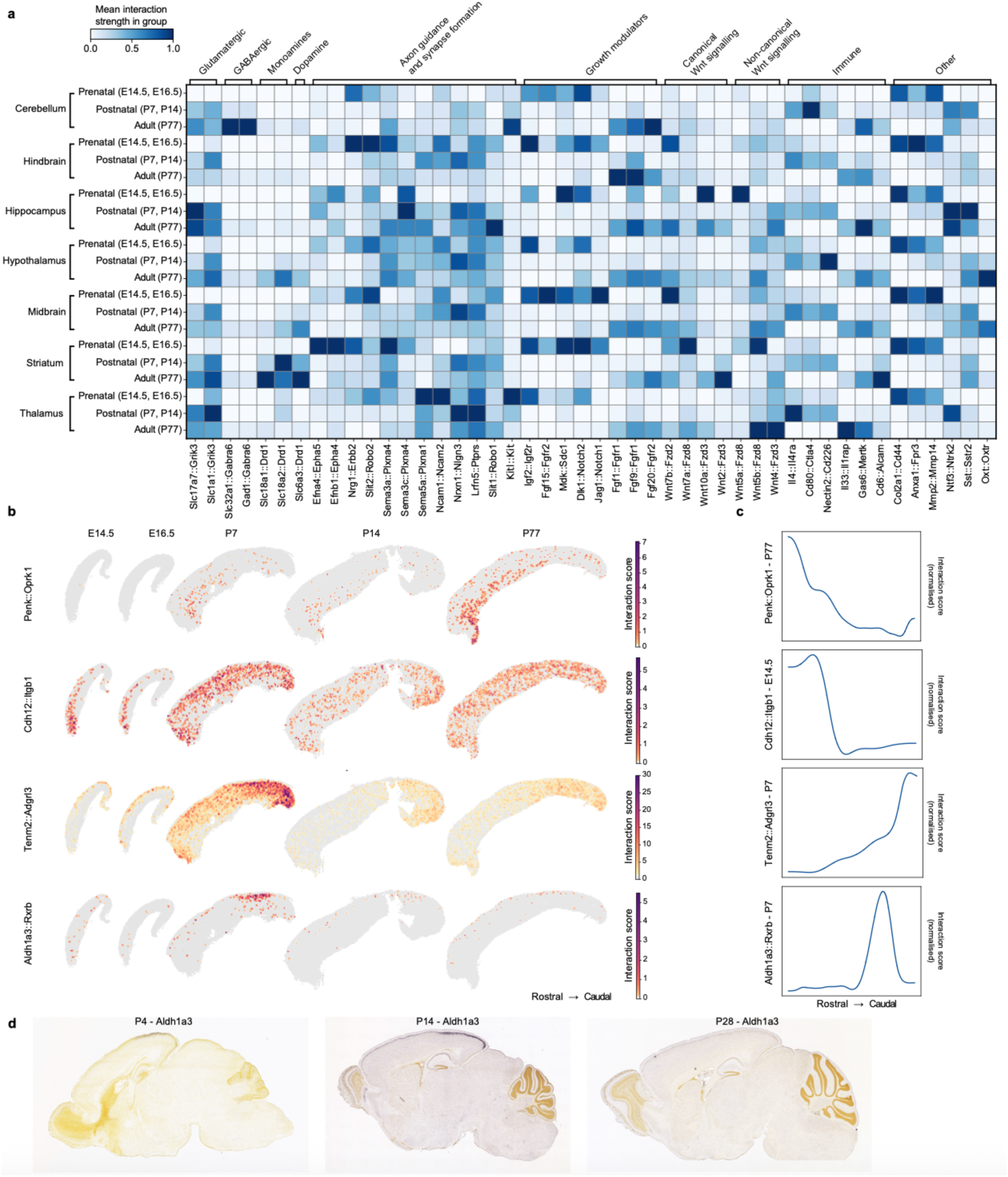
Spatiotemporal analysis of ligand-receptor interactions in the mouse brain cortex with LARIS. **a)** Interacting LR pairs in the annotated brain regions at the prenatal (E14.5, E16.5), postnatal (P7, P14), and adult (P77) developmental time points. Example interactions are grouped according to their functionality, such as neurotransmission (glutamatergic, GABAergic, monoamines, dopamine), axon guidance and synapse formation, growth modulators, canonical and non-canonical Wnt signalling, immune and other. Interaction score is normalized per interaction. **b)** Selected interactions showing enrichment in the rostral-caudal axis. **c)** Normalised LR interaction scores in the rostral-caudal direction for the Penk::Oprk1 at P77, Cdh12::Itgb1 at E14.5, Tenm2::Adgrl3 at P7, and Aldh1a3::Rxrb at P7. **d)** Representative *in situ* hybridization images for *Aldh1a3* at P4, P14, and P28. Image from Allen developing mouse brain atlas: in situ hybridization (ISH) data^56^.

## Methods

### LARIS model

#### Calculation of the Ligand-Receptor integration scores at single-cell resolution

To quantify, at single-cell (or bead) resolution, each cell’s participation in a given LR interaction, LARIS builds a spatial *k*-NN graph on the coordinates. To account for the diminishing probability of CCC with increasing distance, we apply an exponential decay function to the *k-*NN graph. Specifically, for each pair of neighbouring cells with Euclidean distance d_ij_ the edge weight was defined as:

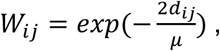

where *μ* denotes the mean of all distances in the graph. This formulation ensures that nearby cells receive weights close to 1, while more distant neighbours are exponentially down-weighted. Gene expression is then “diffused” over space by performing a one-step graph diffusion on the *k*-NN graph:

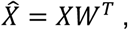

where *X* is the gene expression matrix and *W* is the spatial *k-*NN graph, which naturally aggregates signals from nearby senders and receivers while down-weighting distant cells. To prevent over-emphasizing ligand availability alone, both ligand and receptor expressions are diffused under the same kernel. For each LR (*L*,*R*) pair and cell *c*, we compute an LR integration score as the element-wise product of the diffused profiles:

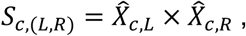

which is high only when both partners are present in the local neighbourhood (e.g., a ligand-expressing cell near receptor-expressing cells). Finally, to avoid scores driven purely by diffusion, we mask out cells that express neither the ligand nor the receptor (i.e., retain scores only where at least one partner is expressed in the cell), yielding spatially grounded LR activity maps whose cell-wise scores can be analysed like pseudo-gene features in downstream visualization and statistics.

#### Identify spatial Ligand-Receptor interaction patterns

To identify LR interactions that exhibit localised spatial structure in a label-free manner, for each interaction we compare the LR integrations scores to a neighbour-smoothed version of the same scores obtained by applying a second diffusion with the spatial *k*-NN graph:

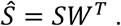

We then quantify their alignment using cosine similarity, which measures the angle between the original and smoothed patterns and is insensitive to overall scale:

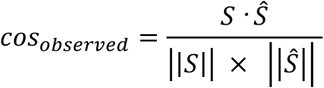

High alignment arises when elevated LR activity concentrates in continuous neighbourhoods (e.g., clusters of nearby high-score cells), whereas diffuse patterns align less strongly. To control for alignment driven by global abundance rather than true specificity, we construct a null by randomly shuffling cell coordinates, rebuilding the *k*-NN graph, repeating the smoothing, and recomputing the cosine similarity. The final spatial-specificity score is calculated by the difference between the observed and shuffled cosine similarity scores:

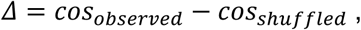

and LR pairs are ranked by Δ to prioritize interactions whose spatial organization exceeds that expected under spatial randomization, enabling fast, label-free pattern discovery.

#### Score interactions per cell type as senders or receivers of communication

For each LR and ordered cell type pair, we assign a directional score that is the product of three components: specificity of the ligand and receptor, abundance of the LR interaction, and neighbourhood enrichment to detect co-localisation. Specificity is calculated using COSG to quantify how specific/characteristic each ligand and receptor is of the putative sender and receiver. We take the normalized COSG marker expression specificity strengths and multiply them to obtain a single specificity term. For cell type A expressing ligand L (sender), and cell type B expressing receptor R (receiver) the specificity is calculated as:

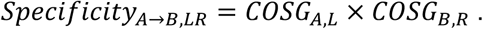

The abundance metric measures the prevalence of LR activity within the cell types: we compute the fraction of cells with a non-zero LR score and combine them using multiplication to penalize cases where one side is rare. For cell type A (sender), cell type B (receiver), and interaction LR the abundance is calculated as:

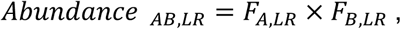

where *F* stands for the fraction of LR score in a cell type.

To quantify co-localisation, we first diffuse the cell type labels over the same spatial *k*-NN graph used above, yielding a set of proximity-to-type scores for every cell: “how enriched is type X (or Y) around this cell”. Next, to characterise cell type pairings, a pairwise coupling matrix (*P*) is constructed by multiplying the proximity-to-type scores resulting in a matrix representing the cell type by cell type combinations. The final metric is then calculated using the cosine similarity between LR integration score and pairwise coupling matrices:

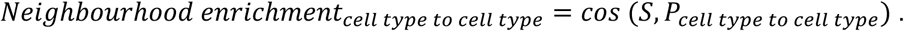

This directly achieves two conditions we want simultaneously: there is LR activity (via the LR integration score matrix), and the putative sender/receiver types are physically proximal (via the diffused labels).

The neighbourhood-enrichment term is multiplied with the abundance and specificity components to yield the final directional score on the cell type level. For cell type A (sender), cell type B (receiver), and interaction LR the final directional score is calculated as:

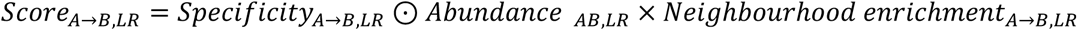

#### Identification of niche enrichment of interactions

To detect LR interactions that are enriched in functional regions or niches independent of cell type, we first assign regional labels to cells (e.g., cortical upper vs. deep layers). Because the LR integration score is defined per cell, standard single-cell differential frameworks can be applied directly: we compare per-cell LR scores across regions to identify interactions that are regionally differentially expressed. In practice, we treat each LR pair as a feature (gene) and perform differential testing across the region groups (e.g., marker-identification/rank-based tests with multiple-testing correction), potentially reporting effect sizes and region-wise prevalence. This yields a list of region-specific LR interactions that reflect spatially localized signaling programs without relying on cell type annotations; if desired, the same analysis can be repeated within cell type strata to differentiate regional effects per cell type.

### Benchmarking performance

#### Simulation of scRNA-seq data

The scRNA-seq data were simulated using a Python implementation of Splatter^10,11^. To mimic a real biological dataset, we used the Tonsil dataset from the Slide-tags study as the reference and fit parameters with Splatter’s splatEstimate, thereby reproducing its dropout rates and expression distributions. We simulated the same number of genes as in the reference transcriptome (26,081) and varied the number of single-cells per simulation according to the associated spatial design. The number of cell types in each simulation was determined by the underlying spatial structure; within a simulation, some clusters were designed to exhibit more distinct differential-expression patterns than others. To avoid unnecessary complexity and potential confounders in benchmarking, we did not simulate batch effects or doublets. Full parameter and design details are provided in Supplementary Table 2.

#### Processing of simulated scRNA-seq data

The simulated raw count matrices were then processed as if they were real data examples using Scanpy^57^. The total unique molecular identifier (UMI) counts were normalised to 10000 UMI per cell using the normalize_total function, then log-transformed with the log1p function, and the top 2,000 highly variable genes were selected with highly_variable_genes. For visualization in two dimensions, we embedded the cells in a UMAP^58^ using the top 50 principal components, all other settings were left at default. Each cell was provided a cluster (cell type) label by the simulation. A simulation was considered successful when the cell types given by the simulation were clearly separated on the UMAP, and when the cell and gene quality metrics matched the tonsil reference (Extended Data Fig. 2).

#### Spatial structure generation

To generate spatial structures, we used two separate approaches. For the fully synthetic structures, scCube’s simulate_cluster and simulate_stripes functions were used^12^. The underlying structure was generated by specifying the shapes, centroids, and sizes of each region and by defining the total cell number to be simulated. The resulting cluster labels with spatial coordinates were then populated with the previously simulated single-cells to create the final spatial data used for benchmarking. For the structures mimicking real tissue examples, the hematoxylin and eosin (H&E) stained samples were manually segmented, separating regions showing different tissue architecture. The segmented regions were then processed in Python by assigning labels to each region, then merging regions based on perceived functionality and tissue morphology. In the final step, the tissue was populated by single-cells from the corresponding simulations. For the examples where H&E staining was not available, the structural schematics were used as the reference structure.

#### Simulation of synthetic interaction database and ground-truth calculations

First, 1,500 genes were randomly designated as ligands and 2,500 as receptors. Ligands and receptors were then randomly paired to produce 4,000 LR interactions, ensuring that every ligand and every receptor appeared in at least one pair; additional pairs were sampled at random until the total reached 4,000. To limit self-interactions at the cell type–cluster level (i.e., cases where both partners act within the same cluster), whenever the most-probable pairing would yield a same-cluster interaction, we re-sampled a new receptor for that ligand.

We reasoned that an ideal candidate for further investigation would be a ligand or receptor that is both specific and abundantly expressed by a given cell type. Thus, our scoring approach focuses on these features. The relevancy score is calculated for each gene in each cluster/cell type, and for gene *i* and cluster *j* as follows:

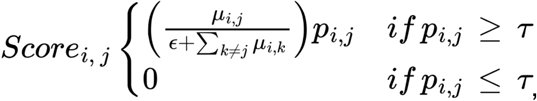

where *μ*_i, j_ is the gene expression of gene *i* in cluster *j*, *p*_i, j_ is the fraction of cells in cluster *j* expressing gene *i*, ɛ is a small constant to avoid division by zero, *μ*_i, k_ is the mean gene expression of gene *i* in cluster *k* (over all cells of type *k*), τ = 0.1: minimum expression-percentage threshold (10%). This scoring identifies the most specific ligands and receptors for each cluster using the simulated parameters.

We generate the ground-truth of interactions at the cluster level. For each potential LR interaction, we calculate an interaction score for each LR pair by multiplying the ligand relevancy score for cluster *i* by the receptor relevancy score for cluster *j*:

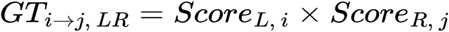

We then factor in spatial information to calculate our spatial ground-truth:

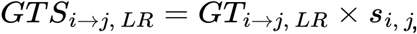

Where *s*_i, j_ denotes the spatial multiplier representing the spatial relationship between cluster *i* and *j*:

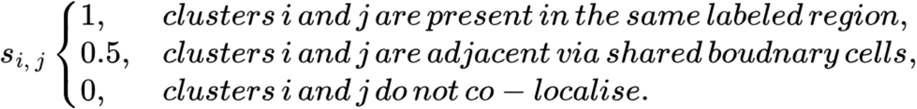

#### CellChat, CellChat spatial, and NICHES analysis

All analyses were run according to the available tutorials. For the CellChat analysis the spatial count matrix was first normalized and log transformed. The spatial conversion factor was set to resemble real life examples with a maximum communication distance of 250 μm, and spot size of 1. The data were then processed using the functions identifyOverExpressedGenes and identifyOverExpressedInteractions with the default parameters. To infer the cluster-level interaction scores the computeCommunProb function was used with trim set to 0.1 in the “truncatedMean” mode. Interactions with less than 10 cells participating were removed with the filterCommunication function. All interactions were treated as ‘Secreted Signaling’. For the NICHES analysis the raw count matrices were SCTransform normalised using the version 5 Seurat implementation^59^, PCA and UMAP were run afterwards using the RunPCA and RunUMAP functions using the top 30 principal components. Before running the prediction the data was imputed using RunALRA. The prediction was run using the RunNICHES function in the “CellToCellSpatial” mode with k=4. The final list of interactions was calculated between all possible cell types. The same LR interaction database was used to benchmark the performance of different methods. Interactions were ranked based on their interaction scores for LARIS, interaction probabilities for CellChat, and Wilcoxon test *P* values and average log2 fold changes for NICHES.

#### Measure performance

To assess the capability of different methods to recover the highest-ranked (and deemed most relevant) interactions in the cluster-level ground truth, we compared the recovery “speed” of each method for the first 100 interactions in the ground truth against the top 1,000 interactions in each method and then calculated the area under the curve (AUC). The proportion of recovered interactions among the top 1,000 ground-truth interactions was calculated by comparing the top 1,000 prediction outputs against the ground truth. To quantify the false-positive rate of the methods, we calculated the rate of interactions between spatially non-proximal clusters, meaning that communication would have been physically hindered in a real-life scenario.

#### Effect of *k-*NN choice and technical reproducibility for LARIS

To measure the effect of the chosen *k-*NN and to demonstrate that LARIS is capable of robust predictions across different spatial technologies, we compared LARIS outputs on four mouse brain hippocampus samples from the Slide-seqV2, Slide-tags, Visium HD, and Xenium datasets. To make the technologies comparable, the gene list was subset to those present in the 5,000-gene Xenium panel. Each sample was processed separately but identically: raw UMI counts were normalized in Scanpy using the normalize_total function and then log-transformed with log1p^57^. LARIS was run with k=20, mu=0.2, sigma=100, and n_repeats=5. Gene modules were identified using the identifyGeneModule function of Emergene with resolution=1.8 ^60^, and four representative modules were selected manually for the Xenium data:Module 1 (’Slc17a7::Gria1’, ‘Slc17a7::Gria2’, ‘Slc17a7::Grin2a’, ‘Slc17a7::Grin1’, ‘Slc1a1::Gria2’, ‘Slc1a1::Gria1’, ‘Slc17a7::Grin2b’, ‘Slc17a7::Gria3’, ‘Slc1a1::Grin2a’, ‘App::Sorl1’, ‘Slc1a1::Grin1’, ‘Slc1a1::Grin2b’), Module 2 (’Sst::Sstr4’, ‘Cadm3::Nectin3’, ‘Lamb1::Sv2a’, ‘Slit2::Robo1’, ‘Efnb1::Epha4’, ‘Efnb2::Epha4’, ‘Slit2::Robo2’, ‘Gdf1::Acvr2a’, ‘Slc32a1::Gabra5’, ‘Ncam1::L1cam’, ‘Cadm1::Nectin3’, ‘Cadm3::Cadm1’, ‘Tenm4::Adgrl1’, ‘Tgfb2::Acvr1b’, ‘Gad2::Gabra5’, ‘Gdf1::Acvr1b’, ‘Tenm4::Adgrl3’, ‘Cntn2::L1cam’, ‘Efna3::Epha3’), Module 3 (’Sema5a::Plxna3’, ‘Efna1::Epha7’, ‘Efna2::Epha7’, ‘Sema5a::Plxna1’, ‘Efna1::Epha4’, ‘Ntf3::Ntrk3’, ‘Tenm1::Adgrl1’, ‘Efna2::Epha4’, ‘Gad1::Gabrd’, ‘Gad1::Gabra2’, ‘Ntf3::Ntrk2’, ‘Inha::Acvr2a’, ‘Gdf10::Acvr2a’, ‘Gdf11::Acvr2a’, ‘Gdf10::Acvr1b’, ‘Tenm1::Adgrl3’, ‘Lamc1::Sv2c’), and Module 4 (’Slc17a7::Grm6’, ‘Thbs4::Itga3’, ‘Dio3::Thra’, ‘Fgf7::Fgfr2’, ‘Slc17a8::Grm2’, ‘Thbs4::Itgb3’, ‘Gas6::Tyro3’, ‘Slc18a3::Chrm1’, ‘Sele::Glg1’, ‘Adcyap1::Adcyap1r1’, ‘Slc1a1::Grm6’, ‘Fgf7::Fgfr1’, ‘Sema3b::Plxna4’, ‘Thbs4::Sdc1’, ‘Gdf2::Bmpr2’, ‘Nts::Sort1’, ‘Sema3f::Plxna4’, ‘Efnb3::Ephb3’, ‘Sema4g::Plxnb2’, ‘Sema3e::Plxnd1’, ‘Sema3a::Plxna1’). To calculate the combined module score for each, we used Scanpy’s score_genes function. To quantify the effect of downsampling of cell numbers and gene counts on *Δ*, the Xenium sample was downsampled using the subsample and downsample_counts functions. The heatmap matrix was created by calculating Pearson’s correlation coefficients between each combination of outputs. The effect of *k-*NN choice was measured on the tonsil dataset by setting various values as *k-*NN (3, 4, 5, 6, 7, 8, 9, 10, 15, 20, 30, 40, 50). The different spatial technologies were compared after running LARIS with knn=20, nu=0.2, sigma=100, and n_repeats=5. The heatmap matrices were created by calculating Pearson’s correlation coefficients between each combination of outputs.

### Data processing

#### Slide-tags human tonsil data

The Slide-tags human tonsil dataset was reprocessed with Scanpy^57^. Cells with fewer than 100 detected genes were removed using filter_cells, and genes expressed in fewer than 3 cells were removed using filter_genes. Cells with total UMI counts greater than 25,000 were also removed to limit the inclusion of doublets. CellBender-filtered raw UMI counts were normalized to 10,000 UMI per cell using normalize_total, then log-transformed with log1p. The top 2,000 highly variable genes were selected with highly_variable_genes. For two-dimensional visualization, we embedded the cells with UMAP using the top 50 principal components; all other parameters were left at their defaults. Cell type annotations were transferred from the original work^2^. The cluster annotated as myeloid cells was reassigned as macrophages, and FDCs were split and annotated as either FDCs or MRCs based on known marker genes.

#### Stereo-seq mouse brain data

The Stereo-seq mouse samples were processed with Scanpy^57^. The original binning of the beads was kept. Whole embryo slices were cropped to keep only the brain. Cells with fewer than 100 detected genes were removed using filter_cells, and genes expressed in fewer than 3 cells were removed using filter_genes. Raw UMI counts were normalized to 10,000 UMI per cell using normalize_total, then log-transformed with log1p. The leiden function was used to cluster the binned cells, and the resulting clusters were annotated as in the original works according to their anatomical regions. If the original annotation was available it was kept, leiden resolution was chosen to achieve clear separation. Upper and deep layers were separated according to the clustering based on known layer specific markers.

### Analysis with LARIS

#### Slide-tags human tonsil data

To calculate the integrated LR scores for each cell, we applied the prepareLRInteraction function from LARIS (number_nearest_neighbors=20). Interactions with spatial specificity and variability, relative to randomly shuffled locations, were identified using the runLARIS function (n_nearest_neighbors=20, mu=0.25, n_repeats=5, sigma=100). To infer and rank interactions at the cluster level, we employed the _calculate_laris_score_by_celltype function (mu=100, mask_threshold=1e-6) within the runLARIS function. To cluster B germinal centre cells by their interactions, we first filtered the cluster-level predictions to the top 3,000 highest-ranked interactions and selected those LR pairs in which B germinal centre cells were either senders or receivers, yielding a total of 161 interactions. The B germinal centre cells were then subset, their LR scores log-transformed, and PCA and UMAP were constructed on the LR scores with default parameters. The leiden function was used to cluster the cells (resolution=1.1), and the resulting clusters were assigned to dark or light zones based on known marker genes. To visualize the DZ (‘CXCR4’, ‘AICDA’, ‘FOXP1’, ‘MME’) and LZ (‘CD83’, ‘LMO2’, ‘BCL2A1’) genes as modules, we used the score_genes function. Regionally enriched interactions were identified with COSG (mu=100, expressed_pct=0.15, remove_lowly_expressed=True, n_genes_user=100)^61^. To reannotate cells previously assigned as FDCs, we combined gene expression data with the LR scores from the top 3000 highest-ranked interactions at the cluster level (168 in total) into a single object. We then constructed PCA (n_comps=10) and UMAP with neighbours (n_pcs=10, n_neighbors=15). The leiden function was used to cluster the cells (resolution=0.5, n_iterations=5), and annotation was performed based on known marker genes. To test for neighbourhood enrichment at the cell type level, for each cell we collected the neighbourhood composition by cell type labels within a 100-micron radius; we then randomly shuffled the cell type labels 100 times and recalculated the composition each time. The final score is the ratio of the real enrichment value to the average of the random neighbourhood compositions, with 1 representing random enrichment, values smaller than 1 indicating depletion, and values larger than 1 indicating enrichment. Gene Ontology Biological Process (GO_Biological_Process_2021) gene set enrichment analysis was performed in Python using GSEApy on the genes with differential LR scores identified by a Wilcoxon rank-sum test (log_2_[FC] > 1 and P_adj_ < 0.05)^41,62^. The LR pairs were split into ligands and receptors before the test.

#### Stereo-seq mouse brain data

To calculate the integrated LR scores for each bin, we applied the prepareLRInteraction function from LARIS (number_nearest_neighbors=10). To identify interactions showing enrichment across different developmental time points and spatial locations, we applied COSG (mu=100, expressed_pct=0.05, remove_lowly_expressed=True, n_genes_user=250) to the regional annotations. To quantify changes in interaction strength in the dorsal-ventral direction, the tissue slices were first oriented to be parallel with the x-axis, then the LR scores were projected into 2D. To quantify changes in the rostral-caudal direction, the tissue slices were first unfolded along the midline, then the LR scores were projected into 2D along the y-axis. Gene set enrichment analysis was performed in Python using GSEApy, with the Kyoto Encyclopedia of Genes and Genomes (KEGG) used as the gene set for the prenatal, postnatal, and adult developmental stages, on the genes with differential LR scores identified by COSG (mu=100, expressed_pct=0.02, remove_lowly_expressed=True, n_genes_user=100). The LR pairs were split into ligands and receptors before the test.

## Data availability

All data analysed in this article are publicly available through online sources. We included links to all data sources in Supplementary Table 3.

## Code availability

LARIS is available as a Python package on https://github.com/genecell/LARIS.

## Acknowledgements

We thank members of the Fishell and Chen labs for comments and discussion. We thank Y. Xu for discussion. This work was supported by grants from the National Institutes of Health (NIH) 5UH3MH120096-05, 1UF1MH130701-01, R01NS081297 and R37MH071679 to G.F., R01HG010647 to F.C., the William Randolph Hearst Fund (FY20) to S.J.W., F32 fellowship from National Institute of Mental Health (NIMH) F32MH125464 to S.J.W. F.C. acknowledges support from the Searle Scholars Award, the Burroughs Wellcome Fund CASI award, the Merkin Institute, Harvard Stem Cell Institute, and the NYSCF. F.C. is an NYSCF Roberston Investigator. D.S. acknowledges support from the NHLBI Pathway to Independence Award (K99HL181185) and the American Lung Association Catalyst Award (CA-1447794).

